# Bacterial survival in microscopic droplets

**DOI:** 10.1101/662437

**Authors:** Maor Grinberg, Tomer Orevi, Shifra Steinberg, Nadav Kashtan

## Abstract

Plant leaves constitute a huge microbial habitat of global importance. How microorganisms survive the dry daytime on leaves and avoid desiccation is not well-understood. There is evidence that microscopic wetness in the form of thin films and micrometer-sized droplets, invisible to the naked eye, persists on leaves during daytime due to deliquescence – the absorption of water until dissolution – of hygroscopic aerosols. Here we study how such microscopic wetness affects cell survival. We show that, on surfaces drying under moderate humidity, stable microdroplets form around bacterial aggregates due to deliquescence and capillary pinning. Notably, droplet-size increases with aggregate-size and the survival of cells is higher the larger the droplet. This phenomenon was observed for 13 different bacterial species, two of which – *Pseudomonas fluorescens* and *P. putida* – were studied in depth. Microdroplet formation around aggregates are likely key to bacterial survival in a variety of unsaturated microbial habitats, including leaf surfaces.

## Introduction

The phyllosphere – the aerial parts of plants – is a vast microbial habitat that is home to diverse microbial communities ^1–6^. These communities, dominated by bacteria, play a major role in the function and health of their host plant, and take part in global biogeochemical cycles ^1–6^. Hydration conditions on plant leaf surfaces vary considerably over the diurnal cycle, typically with wet nights and dry days ^7–10^. An open question is how bacteria survive the dry daytime on leaves and avoid desiccation.

While leaf surfaces may appear to be completely dry during the day, there is increasing evidence that they are frequently covered by thin liquid films or micrometer-sized droplets that are invisible to the naked eye ^11–13^ (Fig. 1A). This microscopic wetness results, in large part, from the deliquescence of hygroscopic particles that absorb moisture until they dissolve in the absorbed water and form a solution. One ubiquitous source of deliquescent compounds on plant leaf surfaces is aerosols ^14–16^. Notably, during the day, the relative humidity (RH) in the boundary layer close to the leaf surface is typically higher than that in the surrounding air, due to transpiration through open stomata. Thus, in many cases the RH is above the deliquescent point, leading to the formation of highly concentrated solutions in the form of thin films (< a few µms) and microscopic droplets ^11^. The phenomenon of deliquescence-associated microscopic wetness is under-studied, and little is known on its impact on microbial ecology of the phyllosphere and on its contribution to desiccation avoidance and cells’ survival during the dry daytime.

**Fig. 1.**
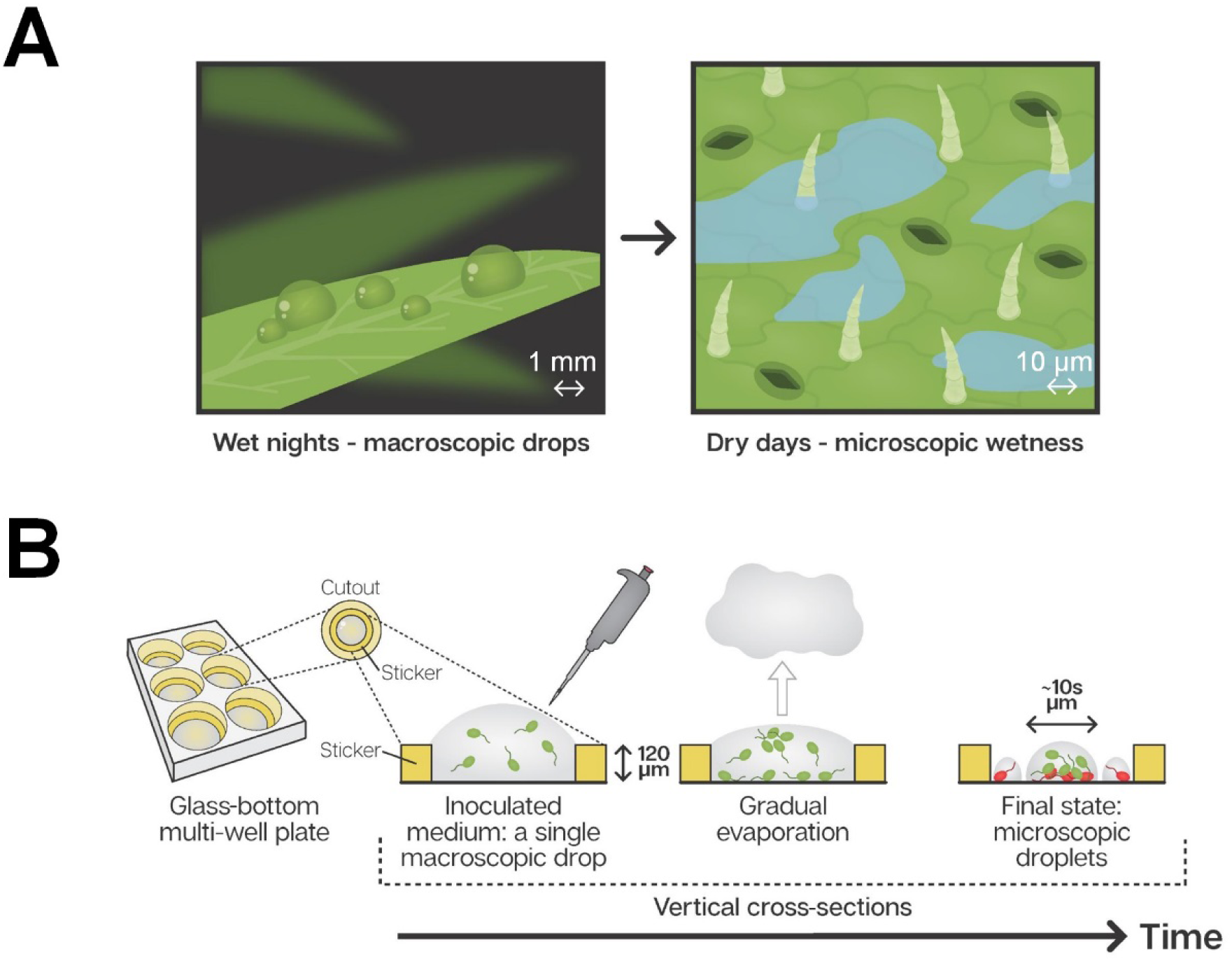
Microscopic wetness: Experimental setup. **(A)** Plant leaf surfaces are usually wet at night with visible macroscopic wetness (e.g., dewdrops). During the day, leaf surfaces are typically dry, with microscopic wetness invisible to the naked eye. (**B**) A thin, round sticker is placed in the center of each well in a glass-bottom, multi-well plate. The hollow part of the sticker is loaded with a medium containing suspended bacteria cells. The well-plate is placed under constant temperature, RH, and air circulation. Gradual evaporation of water from the medium proceeds while bacteria grow, divide, and colonize the surface of the well until the surface becomes macroscopically dry and microscopic wetness forms.

The microscopic hydration conditions around bacterial cells are expected to significantly affect cell survival in the largest terrestrial microbial habitats – soil, root, and leaf surfaces – that experience recurring wet-dry cycles. Only a few studies have attempted to characterize the microscopic hydration conditions surrounding cells on a drying surface under moderate RH and deliquescence’s involvement in this process (a nice exception are the soft liquid-like substances wrapped around cells reported by Mendez-Vilas *et al.* ^17^). Bacterial survival in deliquescent wetness has mainly been studied in extremely dry deserts ^18,19^ and on Mars analog environments ^20,21^. Yet, the interplay between droplet formation, bacterial surface colonization, and survival, has not been studied systematically.

Bacterial cells on leaf surfaces are observed in solitary and aggregated forms. The majority of cells are typically found within surface-attached aggregates, i.e., biofilms ^22,23^. This is consistent with the reported increased survival rate in aggregates under dry conditions on leaves, and poor survival of solitary cells ^24–26^. The conventional explanation for the increased survival in aggregates is the protective role of the extracellular polymeric substances (EPS), a matrix that acts as a hydrogel ^27–30^. Here we ask if aggregation plays additional roles in protection from desiccation. We hypothesize that the resulting microscale hydration conditions around cells on a drying surface depend on the cellular organization (i.e. solitary/aggregated cells and aggregate size) and that the microscale hydration conditions (i.e. droplet size) affect cell survival.

To this end, we designed an experimental system that creates deliquescent microscopic wetness on artificial surfaces. This system conserves some basic important features of natural leaf microscopic wetness while eliminating some of the complexities of studying leaf surfaces directly. This system enabled us to perform a systematic microscopic analysis of the interplay between bacteria’s cellular organization on a surface, microscopic wetness, and cells’ survival on surfaces drying under moderate humidity.

We observed that bacterial cells – aggregates in particular – retained a hydrated micro-environment in the form of stable microscopic droplets (of tens of µm in diameter) while the surface was macroscopically dry. We then quantitatively analyzed the distribution of droplet-size, their correlation with aggregate size and the fraction of live and dead cells in each droplet. The significance of our results is discussed in the context of survival strategies on drying surfaces, microbial ecology of the phyllosphere and possible relevance to other habitats.

## Results

### Drying experiments of bacteria-colonized surfaces

Studying bacteria in microscopic wetness directly on leaves poses a significant technological challenge due to strong auto-fluorescence, surface roughness, and films’ and microdroplets’ transparency. We therefore established a simple experimental system, accessible to microscopy, that enables studying the interplay between bacterial surface colonization, cell’s survival and microscopic wetness on artificial surfaces. This system enables capturing microscopic leaf wetness’s central properties, including contribution of deliquescence, droplets’ persistence, thickness, and patchiness (Fig. 1B) (see Methods). We studied in depth two model bacterial strains – *Pseudomonas fluorescens* A506 (a leaf surface dweller strain ^31,32^) and *P. putida* KT2440 (a soil and root bacterial strain extensively studied under unsaturated hydration conditions ^33–36^). Qualitatively similar results were observed for 16 additional strains (13 bacterial species in total, Table S1). Briefly, bacterial cells were inoculated in diluted M9 minimal media onto hollowed stickers applied to the glass substrate of multi-well plates and placed inside an environmental chamber under constant temperature and RH (28°C; 70% or 85% RH) (Fig. 1B, Methods). Results shown here from 85% RH though 70% RH yielded qualitatively similar results.

### Microscopic droplet formation around bacterial cells and aggregates

At 85% RH, the bulk water evaporated after 14±1h, leaving the glass substrate macroscopically dry (to the naked eye). We then examined the surface of the wells after macroscopic drying was attained (see Methods). Remarkably, the surface was covered by stable microscopic droplets, mainly around bacterial aggregates (Fig. 2A-B). Notably, while solitary cells were surrounded by miniscule droplets (possibly similar to those reported by Mendez-Vilas et al. ^17^), larger aggregates (of ∼ 100 cells) were surrounded by large droplets measuring tens of µms in diameter. Microscopic wetness was preferentially retained around bacterial cells for more than 24h, while uncolonized surface areas were completely dry.

**Fig. 2.**
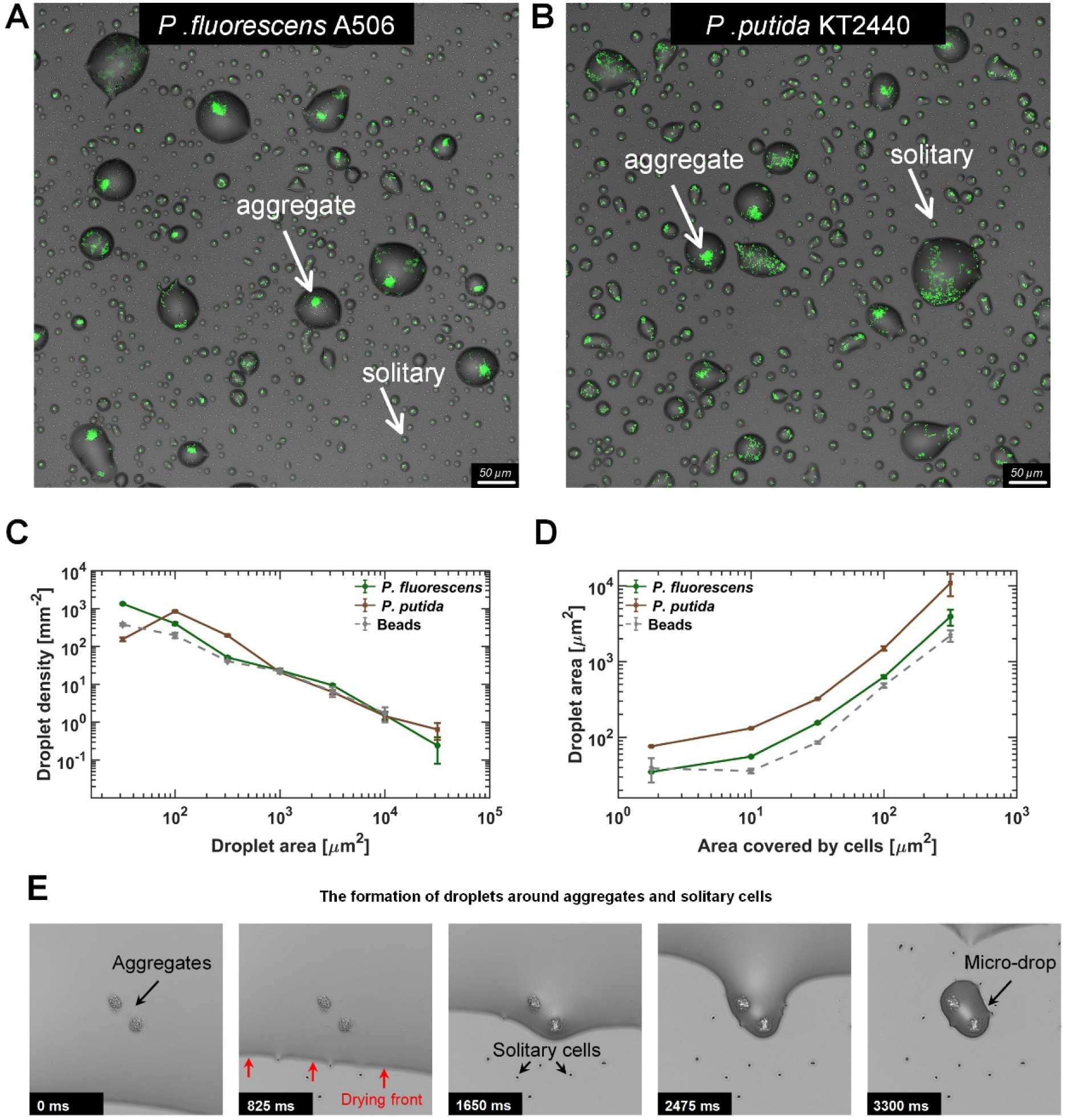
Microdroplets form around bacterial cells and aggregates. **(A-B)** Representative sections of the surface imaged 24h after macroscopically dry conditions were established. Bacterial cells (green) that colonized the surface during the wet phase of the experiment are engulfed by microdroplets, while uncolonized portions of the surface appear to be dry. Solitary cells are engulfed by very small microdroplets, while large aggregates are engulfed by larger droplets (white arrows). Images show a 0.66 × 0.66mm section from an experiment with *P. fluorescens* (A) and *P. putida* (B). **(C)** Droplet-size distributions at 24h: Droplets from both strains show power law distributions with relatively similar exponents (γ = -1.2±0.15 (mean±SEM) and -1.0±0.45 for *P. fluorescens* and *P. putida* respectively). **(D)** Droplet size as a function of cell abundance within the droplet (estimated by area covered by cells): Droplet size increases with cell abundance within drop. Error bars in (C) and (D) are standard errors. (**E**) A time-lapse series capturing the formation of microdroplets around bacterial aggregates: The thin (a few µms) liquid receding front clears out from the surface, leaving behind microdroplets whenever it encounters bacterial cells or aggregates (see also Movie S1).

In order to assess the distribution of droplet size and the correlation between droplet size and aggregate size, we scanned a large area of the surface (∼ 10 mm^2^) to collect and analyze information on thousands of microdroplets (Methods). We found that droplet size (measured by droplet area) follows a power law distribution with similar exponents for the two studied strains (Fig. 2C). When droplet size was plotted as a function of area covered by cells within each droplet (as proxy for cell number, see Methods), a clear positive correlation between cell abundance and droplet size emerged (Fig. 2D).

### The underlying mechanisms of droplet formation

To understand how these microdroplets form, we tested what components of the system were essential for this process. First, we repeated the experiments with fluorescent beads (2µm in diameter) instead of bacteria. Interestingly, we found that microdroplets formed even around beads (Fig. S1), with a similar droplet-size distribution as in experiments with bacteria; and a surprisingly similar correlation between the size of the droplet and the number of beads therein (Fig. 2C-D). In a control experiment without any particulates – bacterial cells or beads – a much smaller number of droplets formed (<1 droplets of >10µm^2^ area per mm^2^, as opposed to >100 droplets of that size in experiments with bacteria). These results indicate that the presence of particles is necessary for droplet formation, whereas biological activity is not. Last, we repeated the beads experiment with pure water instead of M9 medium. This time we did not observe any droplets, indicating that the solutes control droplet formation and retention through deliquescence.

To observe the surface’s final drying phase, we used time-lapse imaging, enabling us to capture the receding front of the remaining thin liquid layer and the formation of microdroplets (Fig. 2E, Video S1). Retention of droplets around aggregates as well as solitary cells, through pinning of the liquid-air interface (due to the extra capillary force exerted by the presence of roughness ^37^, here in the form of particulates), is clearly evident (Fig. 2E). This phenomenon supports the notion that aggregates’ sizes (but possibly also other properties) determine droplet size. In summary, both deliquescence and capillary pinning are essential components for the differential formation and retention of microscopic wetness around cells and aggregates.

### Cell survival rate increases with droplet size

As cells inhabit a heterogeneous landscape of droplets of various sizes, we next asked whether droplets’ sizes impact cell survival. We applied a standard bacterial viability assay by adding propidium iodide (PI) to the medium (see Methods). Thus, live cells emit green-yellow fluorescence, while dead cells exhibit red emission (Fig. 3A). The assay’s validity was further confirmed by the observation that following further incubation at 95% RH, YFP-expressing cells were dividing (some were even motile), while red cells lacked signs of physiological activity (Fig. S2, Video S2). Notably, although the overall population distribution along droplet size was strain specific, survival of cells was nearly exclusively restricted to large droplets for both strains (>10^3^µm^2^ area; Fig. 3B-C, Methods). *P. putida* showed higher overall survival than did *P. fluorescens* (16% vs. 7%, 24h after drying). We note that the overall survival often varied between experiments, and in some cases *P. fluorescens* had higher survival than *P. putida*. Importantly, regardless of this stochasticity, common to all experiments was a clear trend for both strains: The fraction of live cells within droplets increases with droplet size (Fig. 3D).

**Fig. 3.**
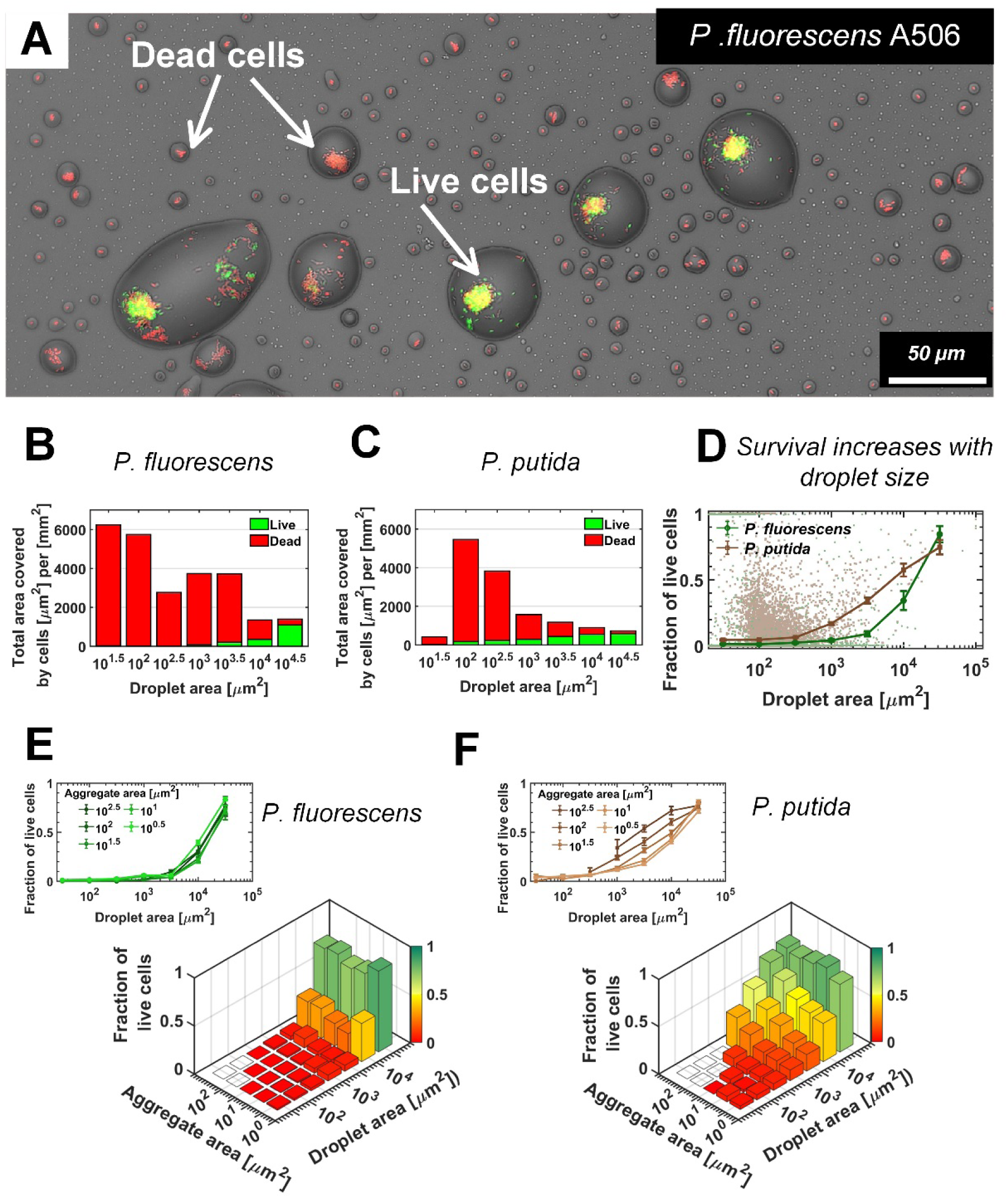
Bacterial survival increases with droplet size. **(A)** A section of the surface covered with droplets (experiment with *P. fluorescens,* 24h after macroscopic drying): Live cells are green, and dead cells (cells with damaged membrane) are red. Live cells were mostly observed in large droplets. **(B-C)** *P. fluorescens* (B) and *P. putida* (C) cell distributions, binned by droplet size: The green and red colored bars indicate the fraction of live and dead cells respectively. **(D)** Fraction of live cells as a function of droplet size. Survival rate increases with droplet size in both studied strains. Error bars represent standard errors. Each dot in the background represents a single droplet. **(E)** *P. fluorescens* survival rates as a function of aggregate size and droplet size: The height of the bars indicates all cellular-objects’ (aggregates or solitary cells) mean survival rates within a given bin of aggregate and droplet size. The inset above shows the same data, but presented differently, with each line representing an aggregate-size bin. Note that there is no pronounced difference between lines, indicating that aggregate size has only minor effect on *P. fluorescens* survival. **(F)** Same as (E) but for *P. putida*: Note there is a pronounced difference between lines, indicating that aggregate size contributes to *P. putida*’s survival (larger aggregates have higher survival); yet droplet size contributes to survival more profoundly than does aggregation (see also Fig. S4).

Accordingly, survival probabilities in small droplets (<10^2^ µm^2^ area) were poor (<5%), in contrast to > 50% survival of both strain*s* in the largest droplets (>10^4^ µm^2^).

Next, we sought to understand what is the net contribution of droplet size to survival. Analysis of cell survival rates as a function of both aggregate size (which by itself affects survival ^24^, cf. Fig. S3) and the size of the droplet they inhabit, shows that for both strains, droplet size strongly affects survival, whereas aggregate size has only marginal (*P. fluorescens*) or moderate (*P. putida*) effect on survival (Fig. 3E,F). Each of these two variables’ relative contributions was also assessed by a multinomial logistic regression model, giving significantly higher weight to droplet size in comparison to aggregate size, for both strains (Fig. S4).

To further study droplet size’s effect on survival, we repeated the drying experiment, but inoculated the cells at a later stage, closer to the macroscopic drying stage, so that the cells did not have time to form aggregates, and were thus mostly solitary. Notably, live cells were observed nearly exclusively in large droplets (>10^3^ µm^2^ area, cf. Fig. S5), and survival increased with droplet size. These results indicate that large droplets promote cell survival. Experiments with 16 additional strains, in both solitary and aggregated cells, yielded qualitatively similar results to those described in the preceding paragraphs (TableS1).

### Formation of droplets using dissolved solutes and microbiota from natural leaves

Lastly, we repeated our experiments using solutes and microbiota extracted from the surface of a natural leaf. We found that stable microdroplets also formed around natural microbiota cells, in some cases only at lower temperatures, suggesting that condensation is involved in microdroplet formation. Furthermore, the microscopic wetness on natural leaf wash was visibly similar to those in our experiments with inoculated bacteria and a synthetic medium (Fig. S6). Experiments using hydrophobic polystyrene substrate rather than glass also yielded qualitatively similar results (Fig.S7).

## Discussion

Our study demonstrates that cell organization on a surface strongly affects the microscopic hydration conditions around cells. We reveal an additional function of bacterial aggregation: improving hydration by retaining large stable droplets (> tens of µms in diameter) around aggregates, and we show that droplet size strongly affects cell survival. Why survival is enhanced in larger droplets remains an open question. We hypothesize that higher water potential in larger droplets provides favorable conditions; further research is required to test this hypothesis.

We suggest that bacterial self-organization on a surface can improve survival in environments with recurrent drying that lead to microscopic wetness. A simple conceptual model that captures the system’s main components and their interactions is depicted in Fig. 4A. Aggregation is an important feature that can affect self-organization, and, in turn, the resulting waterscape, by increasing the fraction of the population that ends up in large droplets.

**Fig. 4.**
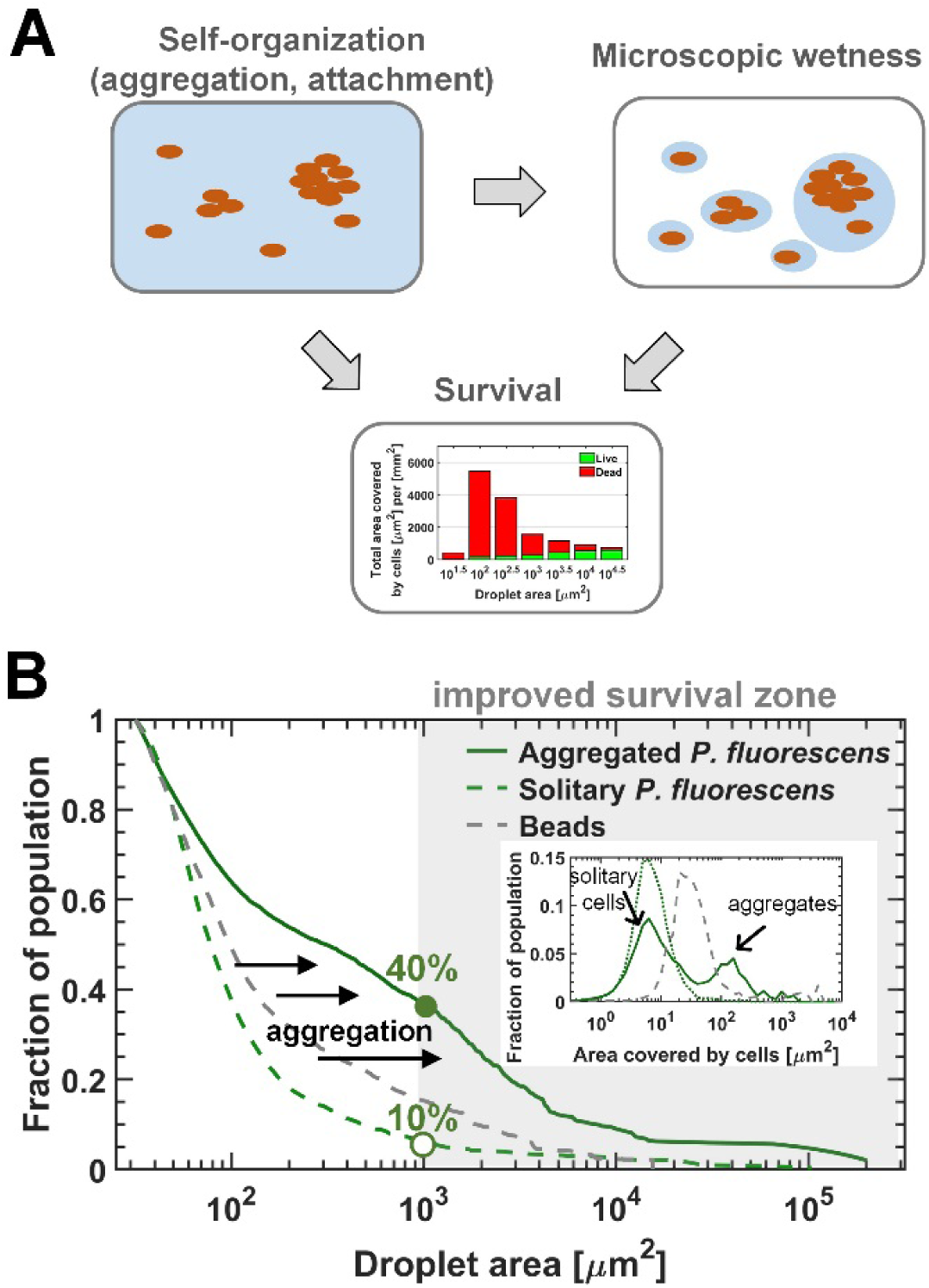
The interplay between self-organization, waterscape, and survival. **(A)** Suggested conceptual model: Cells’ self-organization on the surface affects the microscopic waterscape and the microscopic hydration conditions around cells, which in turn, together with cellular organization (i.e., aggregation) affects survival. **(B)** The three lines represent the fraction of the population residing above a given droplet size of the solitary (late-inoculation) experiment, the bead experiment, and the standard “aggregated” experiment on *P. fluorescens*. Inset: Aggregate-size distributions of these three experiments. Aggregation results in a larger fraction of the population ending up in large droplets with increased survival rates for cells therein.

Preliminary evidence for this is provided by the comparison of the fraction of the population residing in droplets above a given size, using beads, ‘solitary’ and ‘aggregated’ cells as particles (Figs. 4B, S8). The interplay between self-organization, waterscape, and survival is an intriguing open question that merits further research.

Our results suggest that microscopic deliquescent wetness, predicted to occur globally on plant leaves ^11^, can explain how microorganisms survive on leaf surfaces during daytime by avoiding complete desiccation. Yet, they also imply that phyllospheric bacteria have evolved mechanisms to cope with the highly concentrated solutions associated with deliquescent wetness. The ability to tolerate periods of such high salinities could thus be a ubiquitous and necessary trait for phyllospheric bacteria. Better understanding of bacterial survival in microscopic deliquescent wetness, and how it is affected by agricultural practices and anthropogenic aerosol emissions, is thus of great importance to microbial ecology of the phyllosphere and to plant pathology.

Finally, as deliquescent substances are prevalent in many other microbial habitats, it is safe to assume that microscopic deliquescent wetness occurs in many microbial habitats, including soil and rock surfaces ^18,19^, the built environment, human and animal skin, and even extraterrestrial systems (e.g., Mars ^20,21^). Moreover, microscopic wetness is likely to have a significant impact not only on survival, but also on additional key aspects of bacterial life, including motility, communication, competition, interactions, and exchange of genetic material, as demonstrated for soil and other porous media ^28,38^. Microbial life in deliquescence-associated microscopic wetness remains to be further explored.

## Methods

### Experimental design

A simple experimental system, accessible to microscopy, that enables studying the interplay between bacterial surface colonization, cell’s survival and microscopic wetness on artificial surfaces was established (see section **Drying surface experiments)** (Fig. 1B). Fluorescently-tagged bacterial are inoculated in liquid media onto hollowed stickers applied to the glass substrate of multi-well plates and placed inside an environmental chamber under constant temperature and RH (Fig. 1B) (see sections **Drying surface experiments** and **Strains and culture condition)**. After macroscopic drying is achieved, plates are examined under the microscope (see section **Microscopy**) and microscopic wetness, bacterial surface colonization and cell survival are analyzed (see sections **Image analysis** and **Statistical analysis)**.

### Drying surface experiments

Imaging spacers (20mm SecureSeal™ SS1X20, Grace Bio-Labs) were used to confine the inoculum on the surface of 6-well glass bottom plates (CellVis) (Fig. 1B). The spacer was used to reduce flow dynamics effects that result in transfer of biomass to the edge of an evaporating body of liquid drops on flat surfaces (e.g., coffee ring effect ^39,40^). Reduction of flow was achieved through a more spatially uniform evaporation rate. The corners of the spacer were cut to fit the well, adhesive liner was removed from one side of the spacer, and the exposed adhesive was applied to the center of the well by applying gentle pressure against the glass using a sterile disposable cell spreader. The upper liner was removed and the hollow of the spacer was loaded with 340 µl of diluted suspended cells (∼2*10^3^ cell/ml) in half-strength M9 medium (with 2mM glucose conc.). For survival assay, propidium iodide (component B, LIVE/DEAD Bac-Light Bacterial Viability Kit, L-7012, Molecular Probes) was added to the starting inoculum to obtain a final concentration of 20nM. In the experiments with fluorescent beads, rhodamine-tagged micro particles (2µm) based on melamine resin were used (Melamine-formaldehyde resin, FLUKA). The plates were placed, with the plastic lead open, on the uppermost shelf of a temperature- and humidity-controlled growth chamber (FitoClima 600 PLH, Aralab). Temperature was set to 28°C, RH to 70% or 85%, and fan speed to 100%. Prior to the microscopy imaging acquisition, ddH_2_O was added to the empty spaces between the wells of the plate, plates were covered with the plastic lid, and the plates’ perimeter was sealed with a stretchable sealing tape to maintain a humid environment (>95% RH).

### Bacterial strains and culture conditions

*Pseudomonas fluorescens* A506 ^31,32^ and *Pseudomonas putida* KT2440 ^33^ (ATCC^®^ 47054^™^) were chromosomally tagged with YFP using the mini-Tn7 system ^41^ (Plasmid pUC18T-mini-Tn7T-Gm-eyfp and pTNS1, Addgene plasmid # 65031, and # 64967 respectively ^42^). Prior to the gradual drying experiments, strains were grown in LB lennox broth (Conda) supplemented with gentamicin 30 µg/ml for 12h (agitation set at 220 rpm; at 28°C). 50 µl of the 12h batch culture was transferred into 3 ml of fresh LB medium, and incubated for an additional 3-6h (until OD reached a value of ∼0.5-0.7). Suspended cells were transferred to half-strength M9 medium supplemented with glucose by a two-step washing protocol (centrifuge at 7,500 rpm for 2 min., and resuspension of the pellet in 500 µl medium). Half-strength M9 medium consisted of 5.64 g M9 Minimal Salts Base x5 (Formedium), 60 mg of MgSO4, and 5.5 mg of CaCl2 per liter of de-ionized water supplemented with 360 mg glucose as a carbon source (final glucose concentration of 2mM). The full list of strains used in this study is given in Table S1.

### Microscopy

Microscopic inspection and image acquisition were performed using an Eclipse Ti-E inverted microscope (Nikon) equipped with 40/0.95 air objective. An LED light source (SOLA SE II, Lumencor) was used for fluorescence excitation. YFP fluorescence was excited with a 470/40 filter, and emission was collected with a T495lpxr dichroic mirror and a 525/50 filter. Propidium iodide fluorescence was excited with a 560/40 filter, and emission was collected with a T585lpxr dichroic mirror and a 630/75 filter (filters and dichroic mirror from Chroma). A motorized encoded scanning stage (Märzhäuser Wetzlar GmbH) was used to collect multiple stage positions. In each well, 5 xy positions were randomly chosen, and 5x5 adjacent fields of view (with a 5% overlap) were scanned. Images were acquired with an SCMOS camera (ZYLA 4.2PLUS, Andor). NIS Elements 5.02 software was used for acquisition and basic image processing.

### Image analysis

The images were exported from NIS Elements as four separate 16-bit grayscale images per image: bright field (BF), YFP fluorescence (green), propidium fluorescence (red), and a shorter wave-length fluorescence that highlights the droplets (blue). Image analysis was performed in MATLAB. The droplets were segmented by processing the blue fluorescence channel. Droplets were segmented by setting thresholds on the image intensity and gradient following Gaussian filtering (the centers of the droplets are brighter than their periphery and background, and the gradient is more pronounced at the periphery). The two resulting masks were combined, and holes in the connected components were removed. Live and dead cells within each droplet were segmented by histogram-based threshold of the green and red fluorescent channels’ respective intensities, producing binary segmentation and live/dead classification of the cells. The segmented droplets’ image was then used to assign cells and aggregates to their ‘host’ droplet, and to quantify the live/dead surface coverage within each droplet and aggregate.

Our analysis relies on the projected 2D features of 3D objects: droplets and bacterial cells and aggregates. Although some information is lost in the projection, it was deemed a necessary tradeoff for the analysis of the large scanned area and the quantity of data involved. We assume that the relationship between droplet area and volume is monotonous, and that the great majority of cellular aggregates are single-layered. To affirm these assumptions, we performed 3D analysis using z-stacking and 3D deconvolution on a small surface area. This analysis verified that our droplet identification and segmentation does not capture flat discolorations as droplets, and that indeed the cells within the droplets are generally arranged in a single layer on the surface, or suspended in the liquid at densities low enough to maintain the validity of 2D projections.

### Statistical Analysis

Data analyses and statistics for experiments with bacterial cells were based on microscopy images of five different surface sections (each of an area of 2.5 mm^2^) per well. Data analyses and statistics for experiments with beads were based on microscopy images of a surface sections of an area of 10 mm^2^. For statistical analysis of mean values and standard errors, droplets and aggregates were binned by their size on a logarithmic scale. In Fig. 2C standard errors are based on the 5 surface sections (of 2.5 mm^2^) per strain (n=5) and 9 different surface sections (of 1.1 mm^2^) for the beads experiment (n=9). In Fig. 2D and Fig. 3D standard errors are calculated for all droplets within each bin (size range of droplets) of the combined data of the 5 surface sections for experiments with bacteria. In Fig 3B, C and Fig. 4B data is combined for all 5 surface sections. In Fig. 3E,F standard errors are calculated for all aggregates within each bin of the combined data of the 5 surface sections.

## Acknowledgements

We thank Y. Helman, Y. Friedman, Y. Hadar, E. Jurkevitch, O. Yarden, R. Holtzman, O.Bäumchen, D. Sher, A. Bren, and S. Itzkovitz for valuable comments and discussions. We thank S. Lindow, L. Eberl, Z. Cardon, D. Minz, O. Bahar, G. Sessa, S. Burdman, Y. helman and Y. Friedman, for kindly providing bacterial strains. This work was supported by a research grant to N.K. from the James S. McDonnell Foundation (Studying Complex Systems Scholar Award, Grant #220020475).

## Supplementary Information

**Fig. S1.**
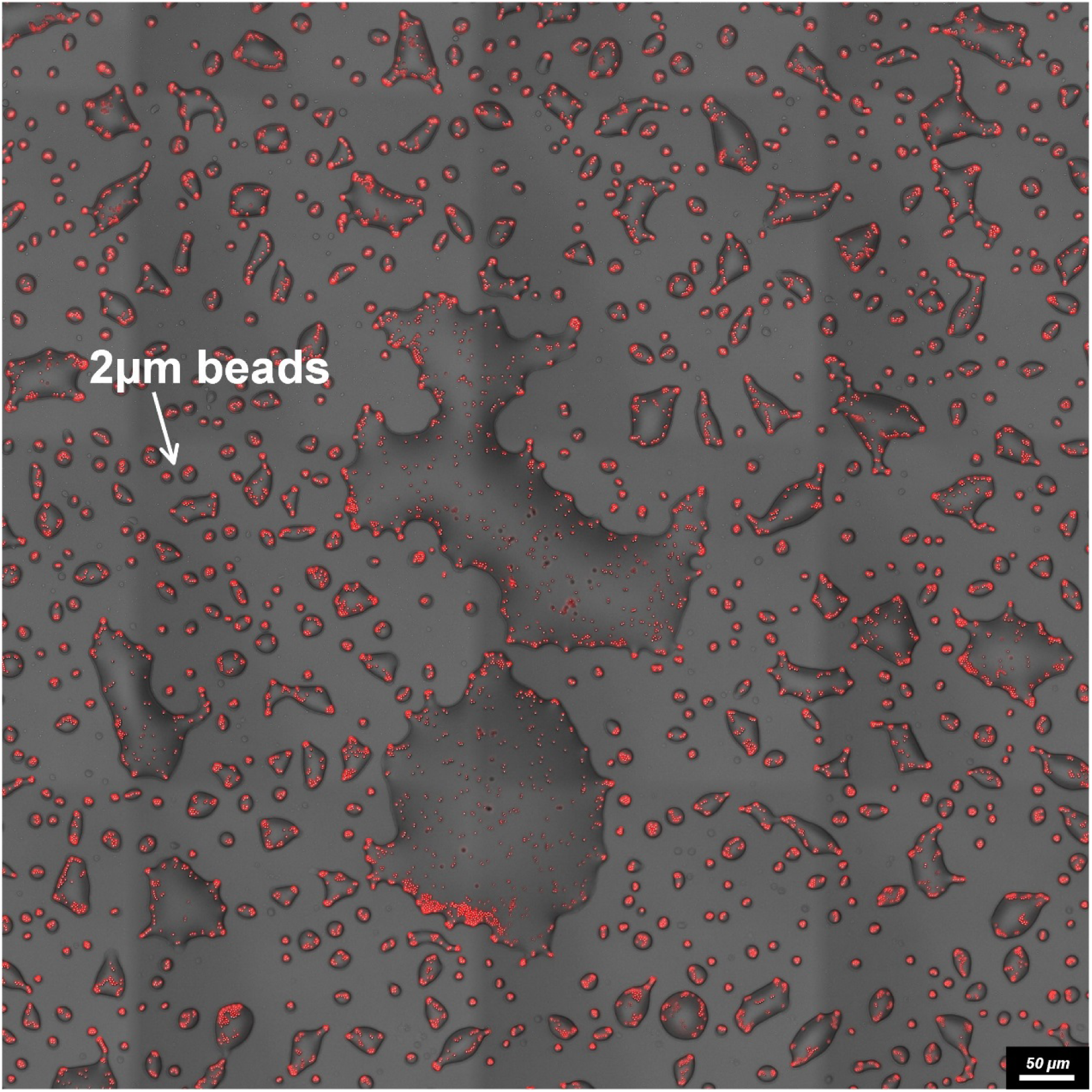
Microscopic wetness – experiment with fluorescent beads (2μm diameter). Experiment was performed under same conditions as the experiments with bacteria (M9 diluted x2, 28°C, 85% RH; see Methods).

**Fig. S2.**
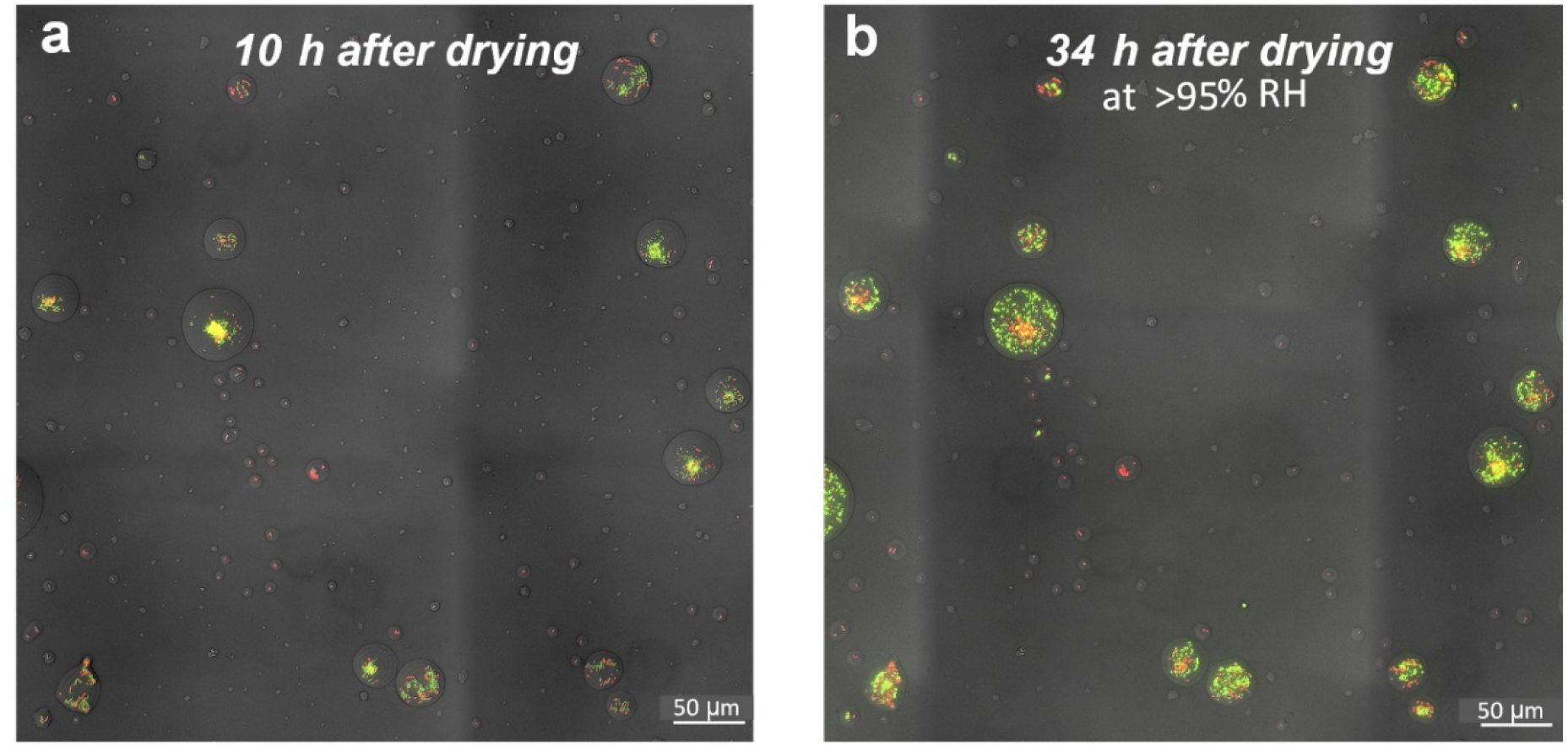
Viability of cells within droplets. **Left:** A section of the surface covered with droplets (experiment with *P. fluorescens,* 10h after macroscopic drying). Live cells are green; dead cells (cells with damaged membrane) are red. Live cells were mostly observed in large droplets. **Right:** Same section of the surface 34h after drying, incubated at ˃95% RH. Red cells in small droplets remain red (dead), while green cells within large droplets disperse (within the droplet boundaries) and divide.

**Fig. S3.**
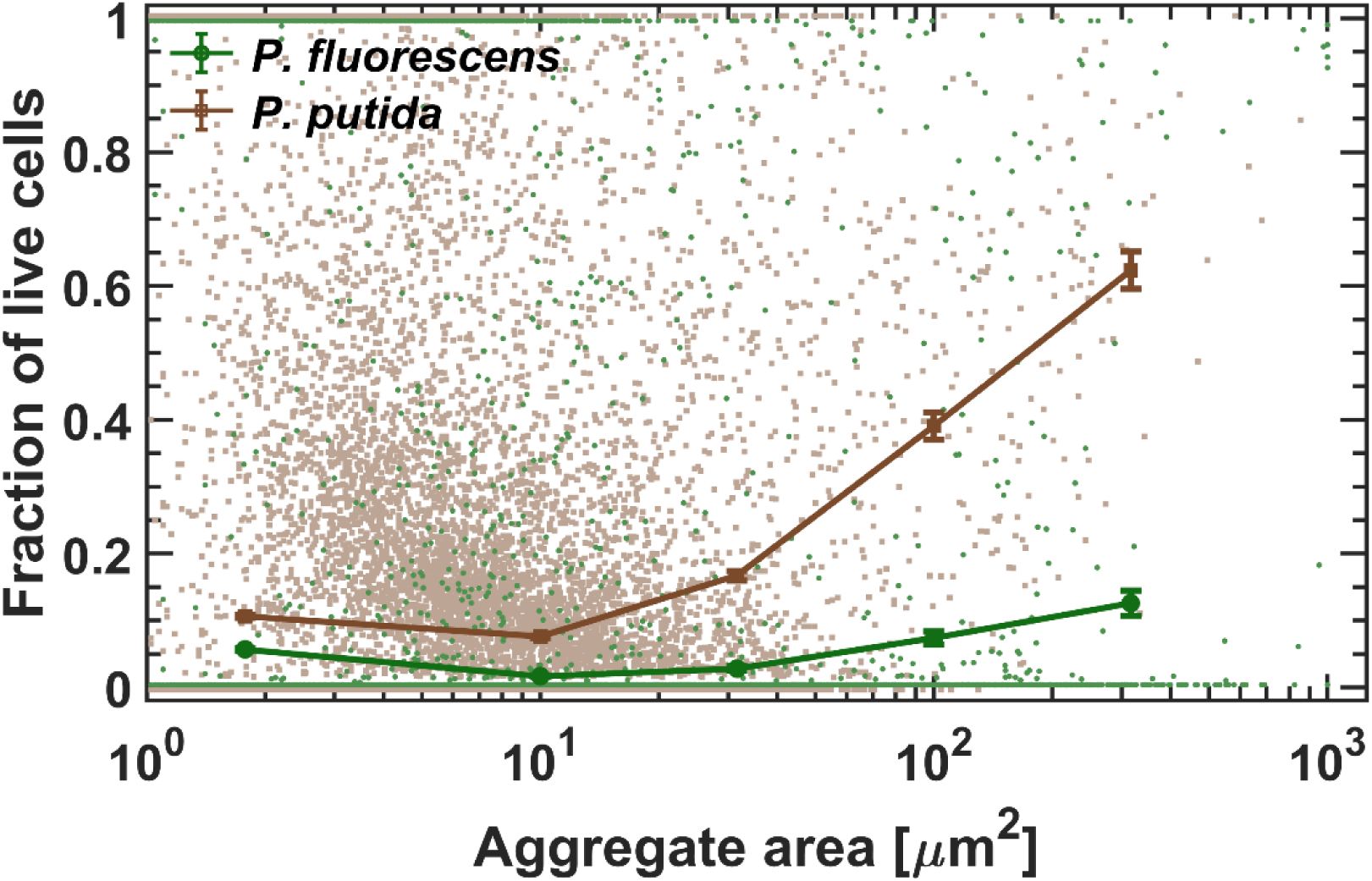
Survival as a function of aggregate size. Survival rate increases with aggregate size in both studied strains. Standard errors are calculated for all aggregates within each bin (size range of aggregates) of the combined data of the 5 surface sections (each of an area=2.5 mm^2^). Each dot in the background represents a single aggregate (solitary cells are aggregates of size ∼1-2 μm^2^).

**Fig. S4.**
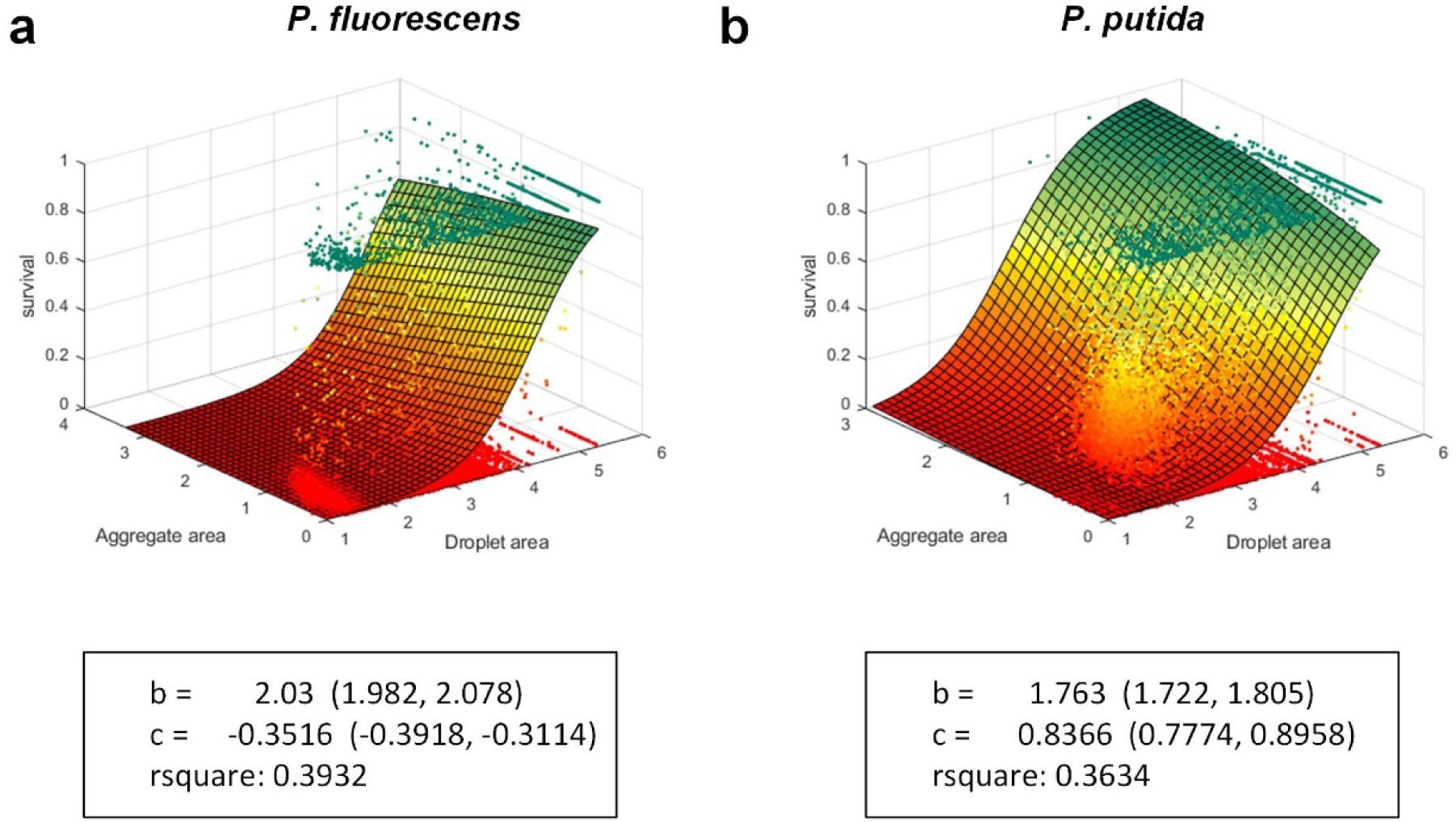
Multinomial logistic regression model. Multinomial logistic regression model was fitted to the data. Each datapoint is an aggregate. The model is defined as z = 1/(1+exp(a - b*x - c*y)) where *x* is log10(host droplet size), *y* is log10(aggregate size), and *z* is survival rate within the aggregate. The fitted points were weighted by the relative area of the aggregate, in order to approximate the distribution of cells. The model is fitted by MATLAB “fit” function. It can be seen via the fitted model outputs that droplet size had a larger coefficient (in absolute values) than did aggregate size for both strains, indicating that droplet size contributes more to survival.

**Fig. S5.**
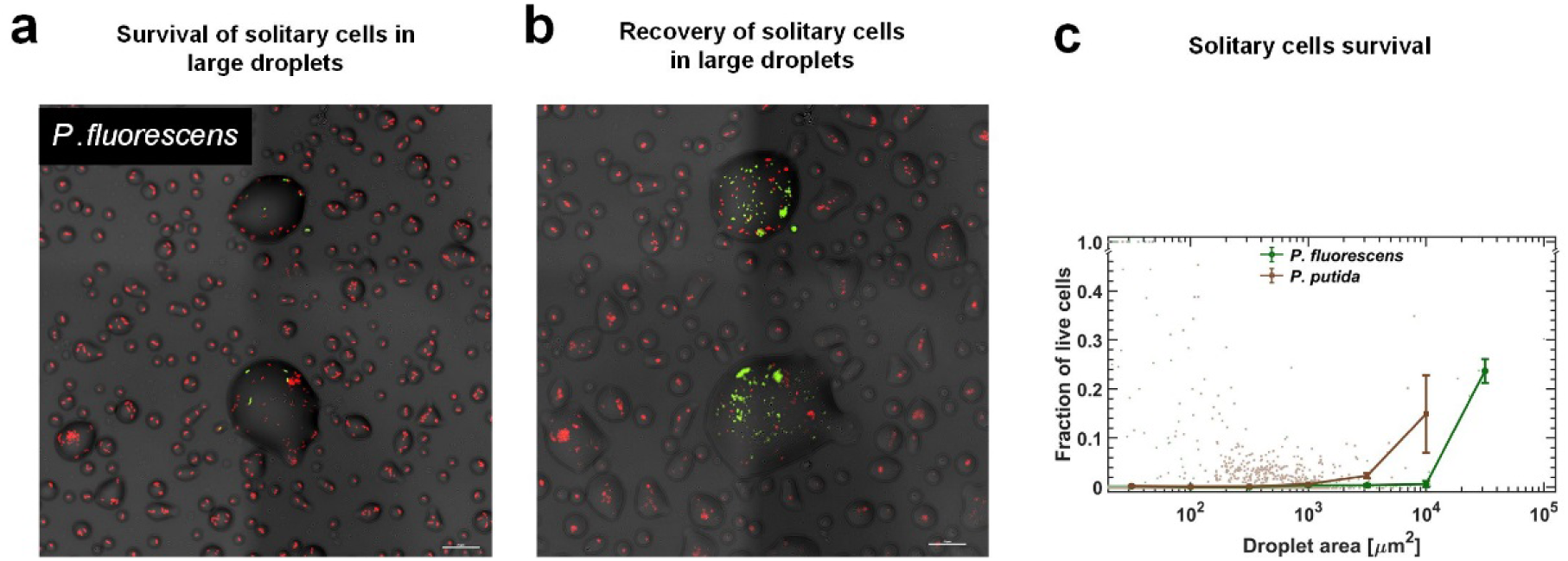
Survival rate of solitary cells increases with droplet size. In this experiment, cells were inoculated at a later stage, close to macroscopic drying (at time = 10h from the beginning of the experiment), and thus did not have enough time to form large aggregates. **(a)** A section of the surface covered with droplets (experiment with *P. fluorescens,* 24h after macroscopic drying). Live cells are green and dead cells (cells with damaged membrane) are red. **(b)** The same section of the surface following another 30h at high RH (∼95%). Recovery of cells was mostly observed in large droplets. **(c)** Survival rate of solitary cells (of both strains) increases with droplet size. Note that overall survival in these experiments was much lower than in the original experiment, pointing to the contribution of aggregation (or self-organization in general) and/or cellular acclimation to overall survival. Standard errors are calculated for all aggregates within each bin of the combined data of the 5 surface sections (each of an area=2.5 mm^2^).

**Fig. S6.**
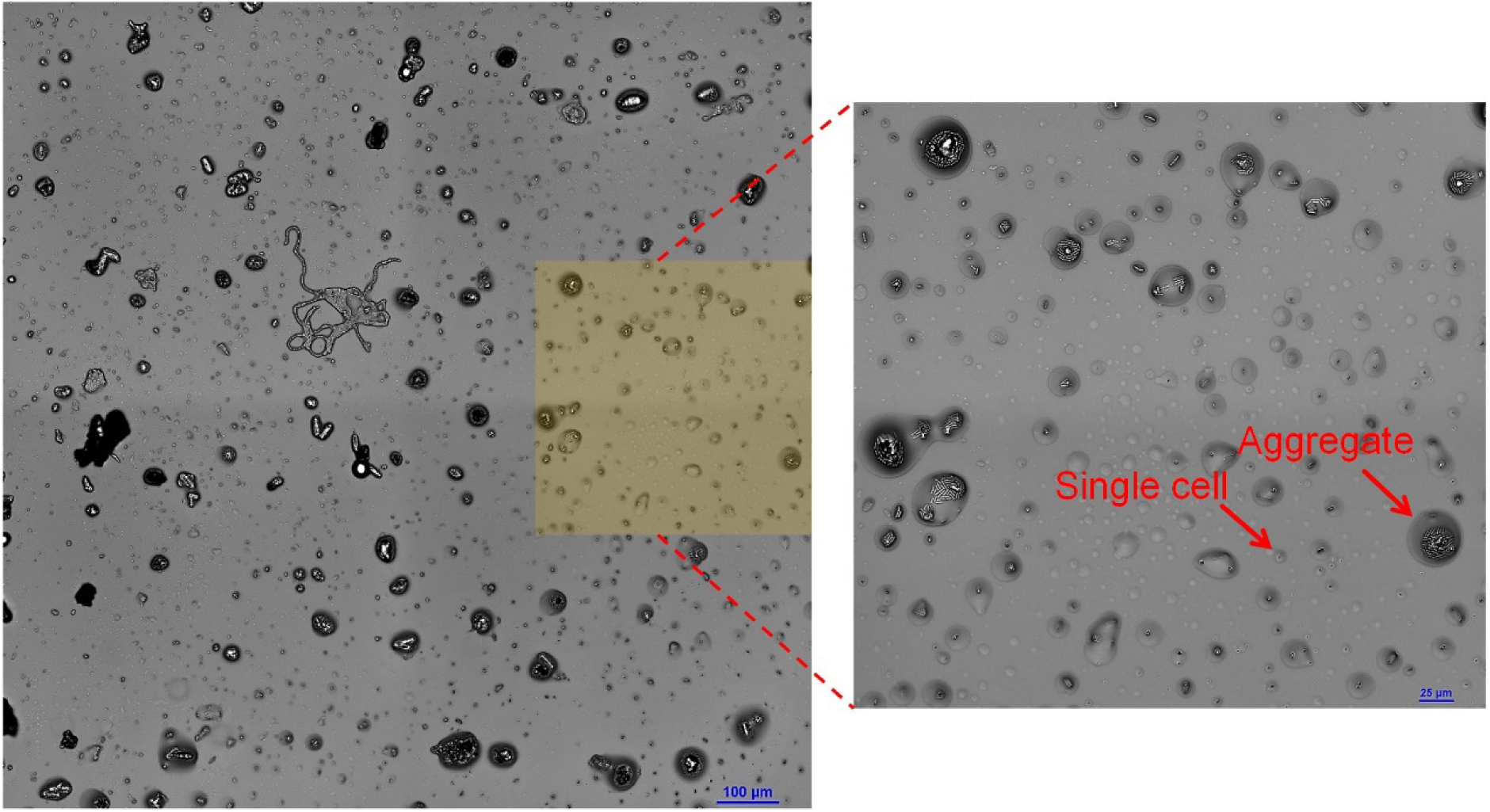
Microscopic wetness forming with natural leaf washes. 340µl of ddH_2_0 were loaded onto the abaxial surface of an ivy leaf, and left for 2h (room temperature). The volume was aspired with a pipette, transferred to our standard gradual drying platform, and left to dry using our described protocol (28°C, 85% RH, see Methods). The plate surface was imaged ∼5 h after macroscopic drying of the well.

**Fig. S7.**
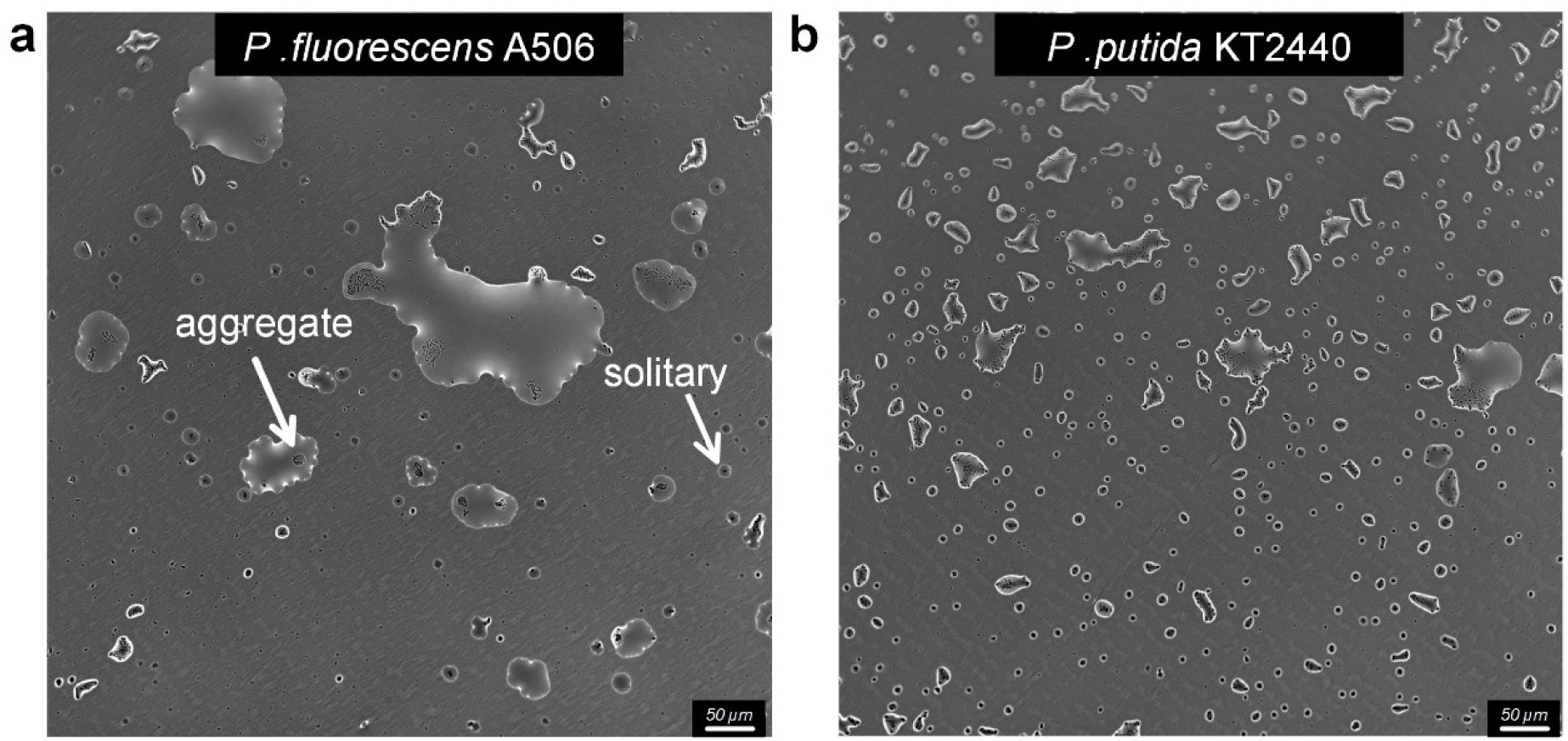
The formation of microdroplets on polystyrene substrate. **(a)** Gradual drying experiment with *P. fluorescens* on a polystyrene 6-well plate (Costar® 6-well Plate, Corning). Representative sections of the surface imaged 48h after macroscopically dry conditions were established. Solitary cells are engulfed by small size droplets while aggregates are found in larger droplets. **(b)** Same as in (a) but with *P. putida*.

**Fig. S8.**
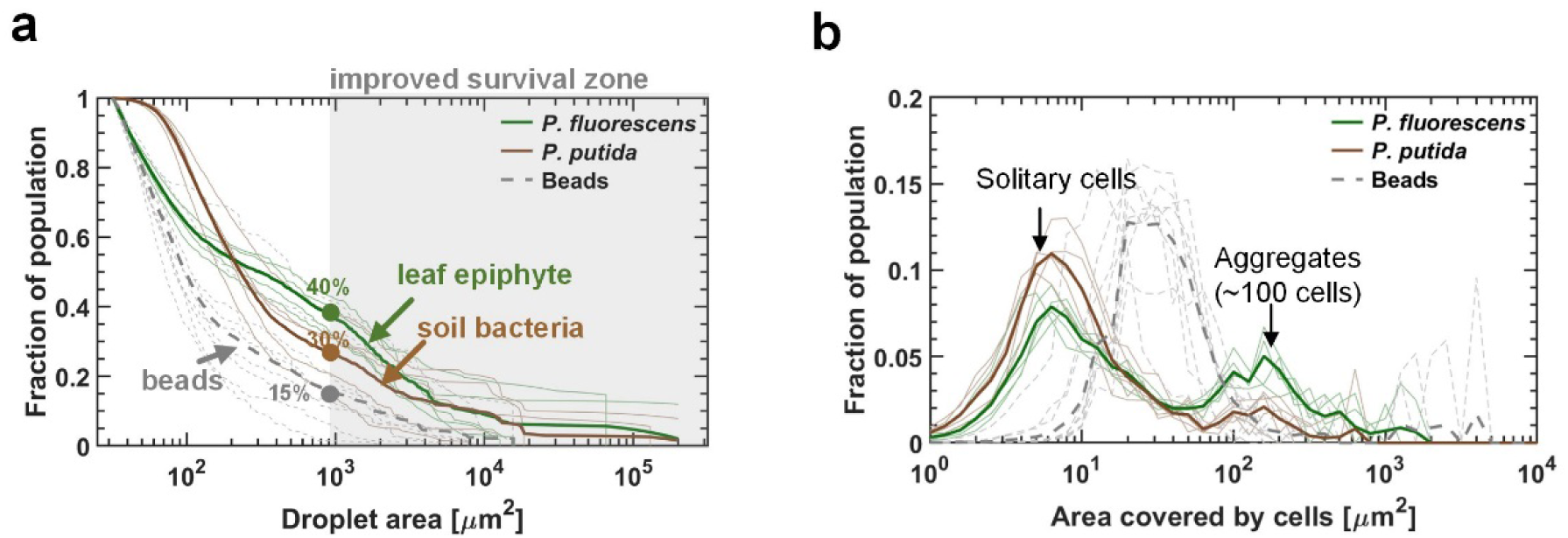
Aggregation and self-organization impact survival. **(a)** The cumulative cell abundance distribution residing above a given droplet size. (**b)** Aggregate-size distributions. In both (a) and (b) the bold lines represent combined data of all 5 surface sections (9 in case of the beads experiment), while the thin lines represent individual surface sections.

**Table S1..** Strains with which the experiments were conducted.

| Genus | Species | Strain | Gram +/- | Major Habitat | Aggregation | Survival at 24h | Gifted from |
| --- | --- | --- | --- | --- | --- | --- | --- |
| *Pseudomonas* | *syringae* | B728a | - | phyllosphere | no | low | S. Lindow |
| *Pseudomonas* | *syringae* | DC3000 | - | phyllosphere | no | low | O. Bahar |
| *Pseudomonas* | *fluorescens* | A506 | - | phyllosphere | yes | medium | S. Lindow |
| *Pseudomonas* | *fluorescens* | NT133 | - | rhizosphere | yes | low | D. Minz |
| *Pseudomonas* | *putida* | KT2440 | - | soil | yes | medium | Purchased from ATCC |
| *Pseudomonas* | *putida* | KT2442 | - | soil | yes | medium | Y. Friedman |
| *Pseudomonas* | *putida* | IsoF101 | - | soil | yes | medium | L. Eberl |
| *Pseudomonas* | *citronellolis* | 13674 (ATCC) | - | soil | yes | low | Y. Friedman |
| *Pseudomonas* | *aurantiaca* | 33663 (ATCC) | - | soil | yes | medium | Y. Friedman |
| *Pseudomonas* | *veronii* | 700474 (ATCC) | - | water | yes | high | Y. Friedman |
| *Pantoea* | *agglomerans* | 299r | - | phyllosphere | no | low | S. Lindow |
| *Pantoea* | *agglomerans* | BRT98 | - | soil | yes | high | Z. Cardon |
| *Escherichia* | *coli* | K-12 MG1655 | - | human gut | no | medium | Y. Helman |
| *Xanthomonas* | *campestris* | 85-10 | - | phyllosphere | no | low | G. Sessa |
| *Burkholderia* | *cenocepacia* | H111 | - | human | yes | low | Y. Helman |
| *Acidovorax* | *citrulli* | M6 | - | phyllosphere | no | low | S. Burdman |
| *Bacillus* | *subtilis* | 3610 | + | soil | no | low | Y. Helman |
| *Clavibacter* | *michiganensis* |  | + | soil | yes | low | S. Burdman |

**Movie S1.**

The formation of microdroplets around bacterial aggregates. The thin (a few *µm*s’ thickness) liquid’s receding front clears out from the surface, leaving behind microdroplets whenever it encounters bacterial cells or aggregates. Video is from an experiment with *P. fluorescens* cells.

**Movie S2.**

Viability of cells within a microdroplet. Some *P. fluorescens* cells can be seen swimming, confined within the droplet.

